# The relationship between pain and depression among middle-aged and older adults in China: Functional limitation as a mediating factor

**DOI:** 10.1101/2025.07.24.666539

**Authors:** Jing Fang, Ningyuan Fang

## Abstract

**Aim:** The association between pain and depression has become a matter of considerable concern in public health. Nevertheless, the fundamental reasons connecting these two phenomena remain unclear. This study sought to explore whether functional limitation serves as an intermediary factor in the link between pain and depression.

**Materials and Methods:** This analysis enrolled 6046 adults aged 45 and older using data from the 2015 wave of the China Longitudinal Study of Health and Retirement (CHARLS)dataset. Statistical approaches included conducting linear regression models alongside bootstrap analyses to evaluate whether functional limitation mediated the association between pain and depression.

**Results:** In models adjusted for confounders, pain intensity showed a positive association with depressive symptoms (β = 0.271, *P* < 0.001) and also exhibited a positive link with functional limitation (β = 0.223, *P* < 0.001). The pathway between pain and depression exhibited a functional limitation that constituted 14.2% of the total effect, with the mediating pathway magnitude estimated at a*b = 0.615(95% CI: 0.521– 0.720).

**Conclusions:** Our findings indicated that pain directly influences depression and also affects them indirectly through functional limitation. Interventions focused on pain management and addressing functional limitation may effectively prevent and alleviate depressive symptoms.

## 1. Introduction

The global population is undergoing an unparalleled demographic shift characterized by an ageing population, with projections indicating that individuals aged>=60 years will constitute a population of 2.1 billion by 2050, representing a critical public health challenge^1^.Concurrently, China is undergoing an accelerated aging process characterized by both scale and velocity unprecedented in modern demographic history^2^. Depressive disorders, operationally defined as psychopathological conditions manifesting persistent affective disturbance, anhedonia, and diminished hedonic capacity^3^, constitute a prevalent neuropsychiatric condition among elderly populations. This disorder is strongly associated with compromised health-related quality of life, with epidemiological evidence demonstrating a positive correlation between age progression and disorder prevalence^4^. Notably, geriatric depression represents a multifactorial public health burden, entailing functional impairment, multimorbidity patterns, heightened healthcare utilization, and elevated mortality risks including suicidality^5^. A recent study analyzing trends in depressive symptoms among middle-aged and elderly individuals in China from 2011 to 2018 found that the prevalence of depression symptoms raised from 36.8% to 44.5% during this period, indicating an increasing trend.^6^ This increase highlights the growing mental health challenges encountered by the elderly population in China, underscoring the need for targeted interventions and enhanced mental health support.

Pain is widely recognized as a significant stressor during the aging process, exerting a profound negative impact on quality of life. It is associated with functional deficits, cognitive decline, and emotional disturbances ^7^. Empirical evidence suggests that individuals experiencing prolonged pain durations exhibit a 154% higher likelihood of developing depression compared to those with shorter pain durations^8^. Chronic pain, in particular, is a major contributor to emotional distress, often culminating in the onset of major depressive disorder (MDD). This association is attributed to the persistent and debilitating nature of chronic pain, which significantly disrupts daily functioning and psychological well-being^9^. Pain and depression are interconnected through shared neurobiological pathways, particularly those involving the neurotransmitters serotonin and norepinephrine^10^. Recent research has identified the paraventricular thalamic nucleus (PVT), specifically its glutamatergic neurons in the anterior PVT, as a critical mediator in the comorbidity of chronic pain and depression^11^. The mechanism linking pain to depression involves neuroimmune interactions, particularly the Corticotropin-Releasing Hormone-Melanocortin-T helper 17 (CRH-MC-Th17) signaling pathway^12^. Chronic pain and depression also are linked through neuroinflammation, characterized by increased levels of cytokines like TNF-a, IL-1b, and IL-6^13^. This inflammatory response exacerbates both conditions. Although many studies have shown great interests in the relationship between pain and depression, the relationships involved are poorly understood. A deeper understanding of these interactions could pave the way for more effective therapeutic strategies targeting both pain and depression, particularly in aging populations.

Functional limitation among the elderly in China represents a significant public health issue, exerting a detrimental impact on cognitive function, healthcare utilization, and overall well-being^14^. A recent study revealed that cancer diagnosis in Chinese adults did not worsen functional limitations; rather, pain may contribute more to functional decline than cancer itself^15^. A longitudinal study conducted among older adults in the United States revealed that individuals experiencing pain exhibited a 52% higher likelihood of developing extremity functional limitations over a six-year period^16^. These findings highlight the importance of pain reduction as a strategic intervention to delay the progression of dysfunction and disability. Furthermore, research indicates that functional limitations serve as a significant mediator linking falls to depressive symptoms in older Chinese adults^17^. Greater functional limitations have been consistently associated with an increased presence of depressive symptoms, emphasizing the interconnectedness of these factors^18^. A recent study demonstrated that pain-related activity interference significantly mediates the association between higher pain intensity and elevated depressive symptoms^19^,which indicated that pain and depressive symptoms may be linked to functional limitations. These findings suggest that functional limitation may be a crucial pathway to mitigating the adverse effects of pain on depression and improving overall quality of life.

In this study, we explored associations between pain and depression, individually or in combination, with functional limitation in middle-aged older people participating in the Chinese CHARLS, and to assess the relationship between pain and depression in Chinese middle-aged and older adults and to examine if functional limitation serves as a mediating role in the aforementioned relationship.

## 2. Patients and methods

The CHARLS is a nationally representative longitudinal survey that is a biennial national study consisting of 4 rounds. As a national study, he surveyed people aged 45 and older and their spouses in China, including an assessment of the social, economic, and health status of community residents. CHARLS examines the health and economic adjustments brought about by the rapid aging of the Chinese population^20^.In our study, we applied CHARLS data collected in the 2015 survey. The baseline sample comprised 21,095 individuals. We excluded participants with incomplete or irregular data values for critical variables, including age (n = 15,049). (Fig 1) Ultimately, the final sample consisted of 6046 participants eligible for inclusion in the analysis.

**Fig. 1.**
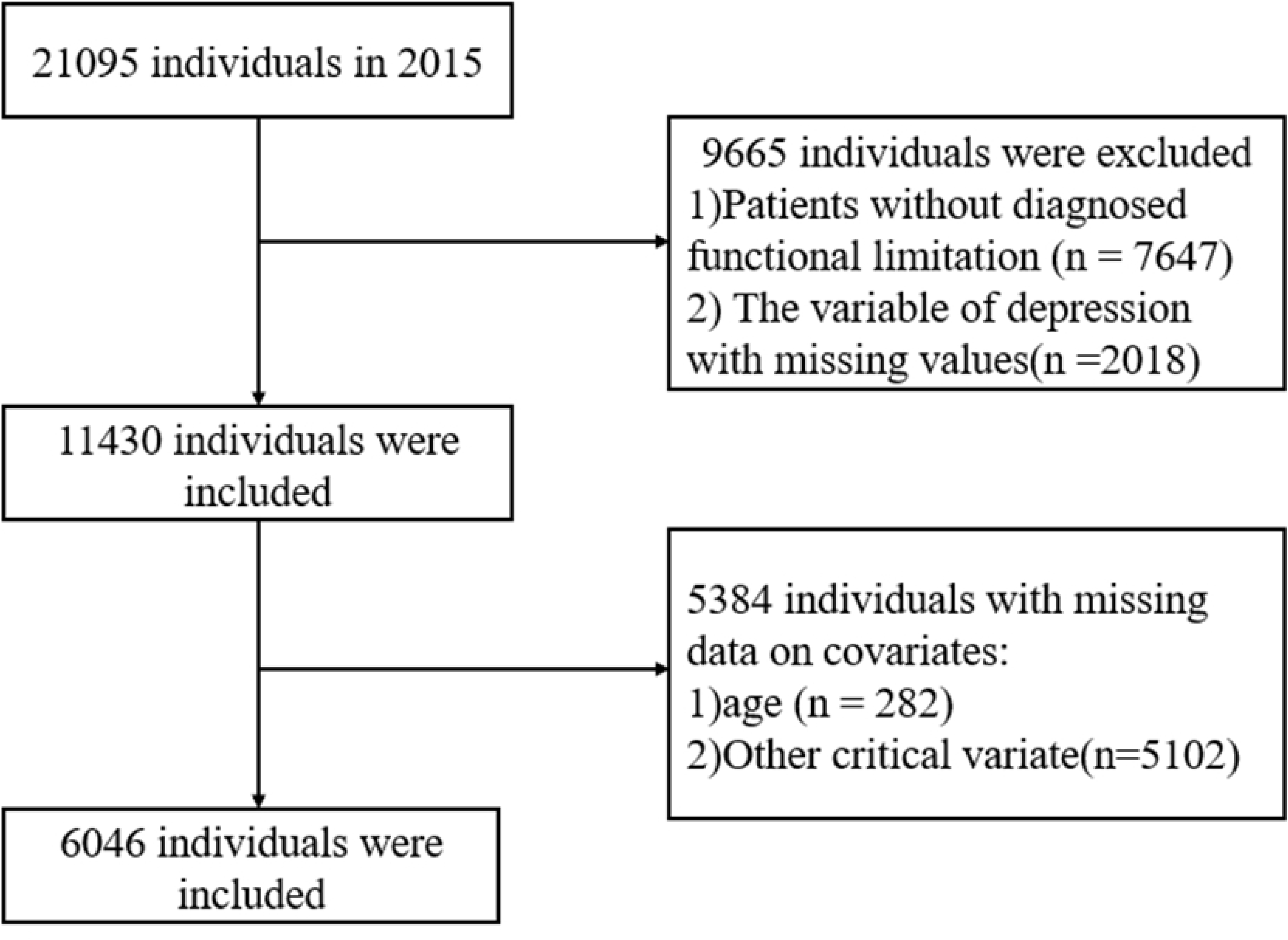
Study Population.

### 2.2. Assessment of Pain

There are 16 types of pain sites in the questionnaire, including head, shoulder, arm, wrist, etc., by asking “What parts of the body do you feel pain?” Please list all parts. Based on this, the presence of pain was defined as self-reported pain in one or more anatomical regions of the body^21^.

### 2.3. Assessment of Depression

It includes 10 areas of the depression as assessed by the Center for Depression 10-item Depression Research Center Depression Scale (CES-D-10)^22^: “I’m bothered by things that don’t normally bother me,” “I’m having trouble focusing on what I’m doing,” “I’m depressed,” “I feel like I’m doing everything I’m working on,” “I’m hopeful about the future,” “I’m feeling scared,” “I’m having trouble sleeping,” “I’m happy,” “I’m feeling lonely,” and “I can’t start.^23^” The options are “Little or no time (< 1 day)”, “Partial or little time (1-2 days)”, “Occasional or moderate time (3-4 days)”, and “Most time (5-7 days)” with a score of 0 ∼ 3, and two questions that reflect positive emotions (I’m hopeful for the future, I’m happy) are implemented in reverse (3 ∼ 0)^24^. The final score is calculated by adding the scores from the 10 questions. The total score ranges from 0 to 30, and when the total score is ≥10, the participant is considered to have depression.

### 2.4. Assessment of Functional limitation

In the 2015 baseline, we used the Instrumental activity of daily living (IADL) and the activity of daily living (ADL) to assess Functional limitation^1^. ADL is defined as challenges in performing basic self-care activities such as dressing, bathing, eating, transferring in and out of bed, using the toilet, and maintaining continence of bladder and bowel. IADL encompasses more complex daily tasks, including housekeeping, meal preparation, shopping, financial management, and medication adherence.^25^. The ADL and IADL answers are on a four-point scale ranging from 1 (no difficulty) to 4 (not possible)^26^. Functional limitations were derived by aggregating ADL and IADL, with total scores spanning from 11 to 44^1^. Higher scores among respondents indicated more severe functional limitation.

### 2.5. Assessment of covariates

Demographic and health-related variables encompassed: sex, age (continuous), educational attainment (categorized as illiterate, elementary, middle school, or high school and above), marital status (married/unmarried), rural/urban residence, smoking/alcohol consumption status (binary), body mass index (BMI, continuous), the number of chronic conditions(continuous), life satisfaction score (ordinal 0-4 scale), and nightly sleep duration (hours). The number of chronic conditions was assessed through self-reported diagnoses of 14 noncommunicable diseases (NCDs), including hypertension, diabetes, chronic pulmonary disease, cardiovascular disorders (e.g., cardiac conditions, stroke), malignancies, musculoskeletal conditions (e.g., arthritis), and others (e.g., dyslipidemia, liver/kidney diseases).Life satisfaction was evaluated via the question^24^: “How satisfied are you with your life overall?” Responses were standardized as:0 = Not at all satisfied; 1 = Not very satisfied; 2 = Somewhat satisfied; 3 = Very satisfied; 4 = Completely satisfied. Scores range from 0 to 4, with higher scores being associated with higher life satisfaction^27^.

### 2.6. Statistical analysis

First, descriptive statistics were calculated to summarize the sample characteristics, including means (standard deviations) for continuous variables and frequencies (percentages) for categorical variables. Next, Spearman correlation tests were conducted to assess the relationships among key variables. For all regression analyses, unstandardized regression coefficients (B) were reported. These coefficients represent the expected change in the dependent variable for a one-unit increase in the independent variable, holding all other variables constant. Standardized regression coefficients (β), also referred to as beta coefficients, were reported separately. The standard errors associated with both unstandardized (B) and standardized (β) coefficients were also reported. Mediation analysis models were used to examine the intermediary role of functional limitations in the association between pain and depressive symptoms among middle-aged and older adults. This involved two steps: (1) a linear regression test to evaluate the direct relationship between pain and depression, and (2) an analysis incorporating functional limitations as a mediating factor. Finally, a non-parametric bootstrap method with 1000 resamples was employed to estimate total, indirect, and direct effects. All analyses were conducted using RStudio 4.3.2 software with a significance level of α= 0.05.

## 3. Results

### 3.1. Basic characteristics

The basic characteristics of the study sample are shown in Table 1 of 6046 individuals, with a mean age of 59.3 years, of which 82.6% (4992) were female. In the sample, about 77.1% of the participants had an education level of high school or above, and the majority of the elderly lived in rural areas (82.2%) and were married (86.5%). At baseline, the mean BMI and number of chronic diseases were 24.5 and 1.5, respectively. In addition, there are significant differences between men and women in a number of ways. For example, the mean age for males was 59.8 years, slightly higher than for females (59.1 years). In terms of educational level, 78.1% of males have a high secondary or higher education, which is higher than that of females (76.9%). Marital status also shows gender differences, with 91.3 %of men married compared to 85.5% of women. In addition, the prevalence of smoking (40.9%) and alcohol consumption (40.5%) among men was significantly higher than that of women (1.8% and 7.8%, respectively). In terms of health-related measures, the average BMI for men was 24.1, slightly lower than 24.6 for women. The average number of chronic diseases was 1.1 for males, significantly lower than 1.5 for females. In addition, 32.7% of men reported pain, lower than 44.0% of women. In terms of psychological and functional measures, the average score of depressive symptoms was 8.0 for men, lower than 9.9 for women, while the mean score for functional limitations was 12.2 for men and slightly lower than 12.6 for women.

**Table 1.**
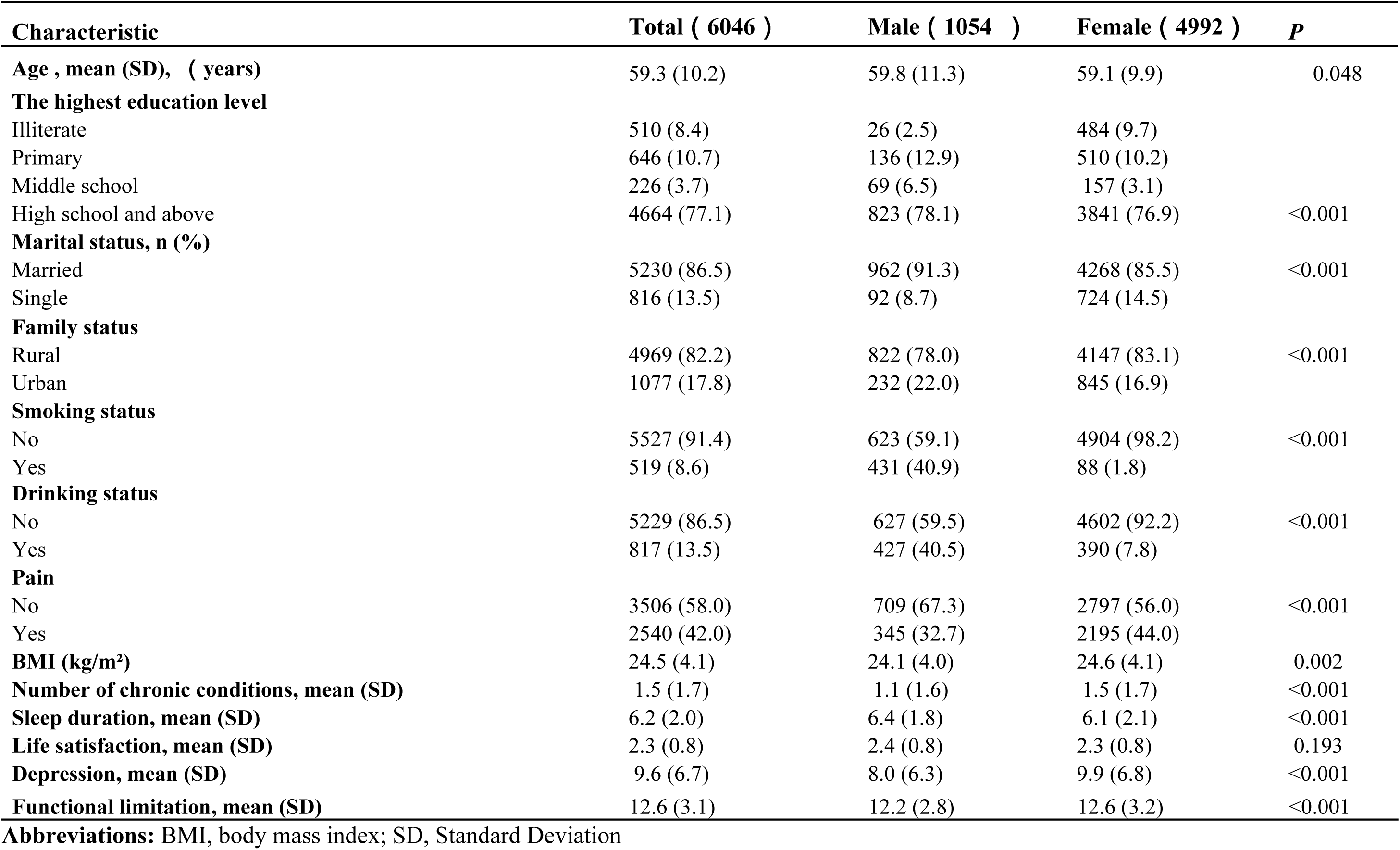
Baseline characteristics of enrolled participants in 2015.

### 3.2. Associations between key variables

Table 2 lists the correlations of baseline pain, functional limitation, and depression. We found that pain at baseline was positively associated with depression (r = 0.45 *p <* 0.001). Pain was positively correlated with functional limitation (r = 0.22, *p* <0.001), and functional limitation was positively correlated with depression (r = 0.35, *p* <0.001). Table2 Correlations of baseline pain, functional limitation, and depression

**Table 2.**
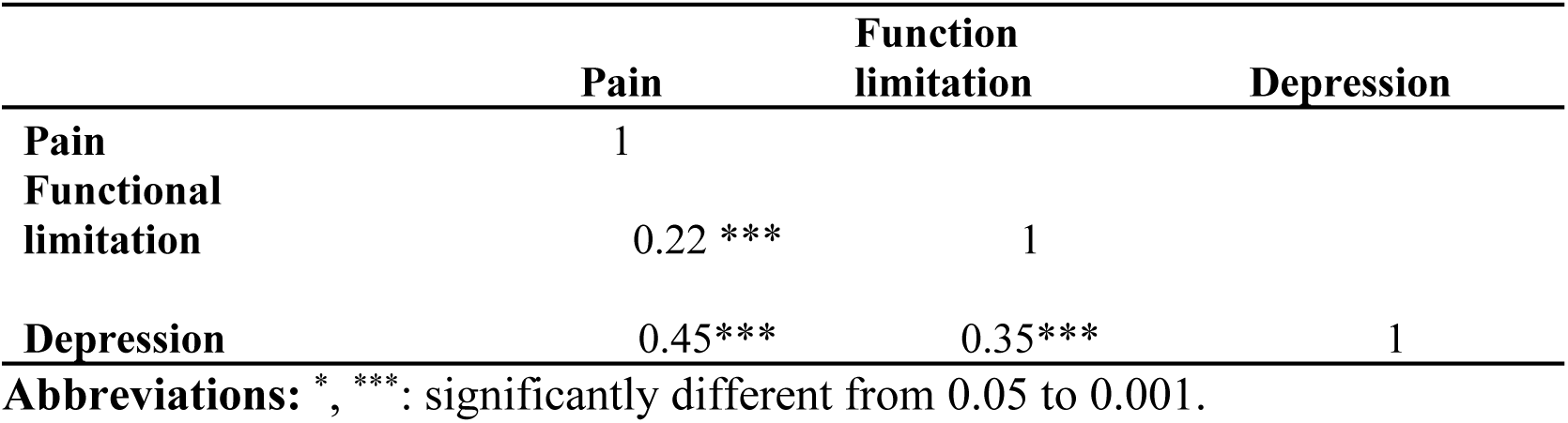
Correlations of baseline pain, functional limitation, and depression.

### 3.3. The mediating role of functional limitation in the relationship between pain and depression

As shown in Table 3, after adjusting for control variables, pain was significantly related with functional limitation (β=0.204, *P <* 0.001). In Table 4, Model 1 indicates that pain is significantly associated with depression (β = 4.311, *P*< 0.001). When functional limitation was incorporated as a mediating variable in Model 2, pain remained significantly associated with depression (β = 3.692, *P*< 0.001). This suggests that functional limitation serves as a mediating factor in the relationship between pain and depression. The reduction in the regression coefficient from Model 1 to Model 2 implies that functional limitation partially accounts for the association between pain and depressive symptoms. However, the persistent significance of pain in Model 2 also indicates that other pathways or factors may contribute to this relationship beyond functional limitation. Therefore, while functional limitation plays a mediating role, it does not fully explain the connection between pain and depression. The results also suggested that functional limitation was a significant mediator of the association between pain and depression, and explained 14.2 % variance of the total effect. The mediation pathway model was shown in Fig 2. The participants were also categorized into different subgroups, including male and female, middle-aged and older adults, and separate analyses were conducted for each subgroup. In all cases, it was demonstrated that functional limitation was a significant mediator of the association between pain and depression (see Supplementary Table S1-S4).

**Fig 2.**
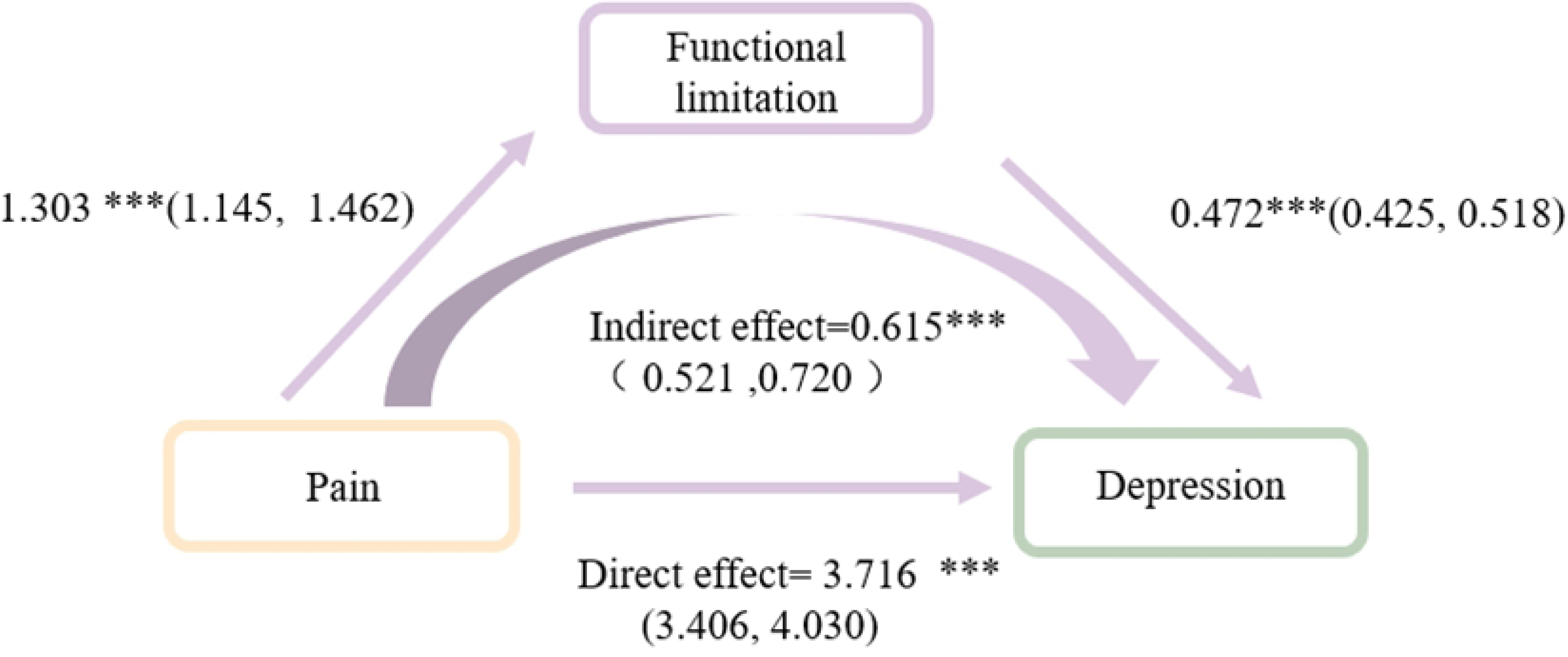
Mediation model visualizing the relationship between pain and depression via functional limitation as a mediator.

**Table 3.**
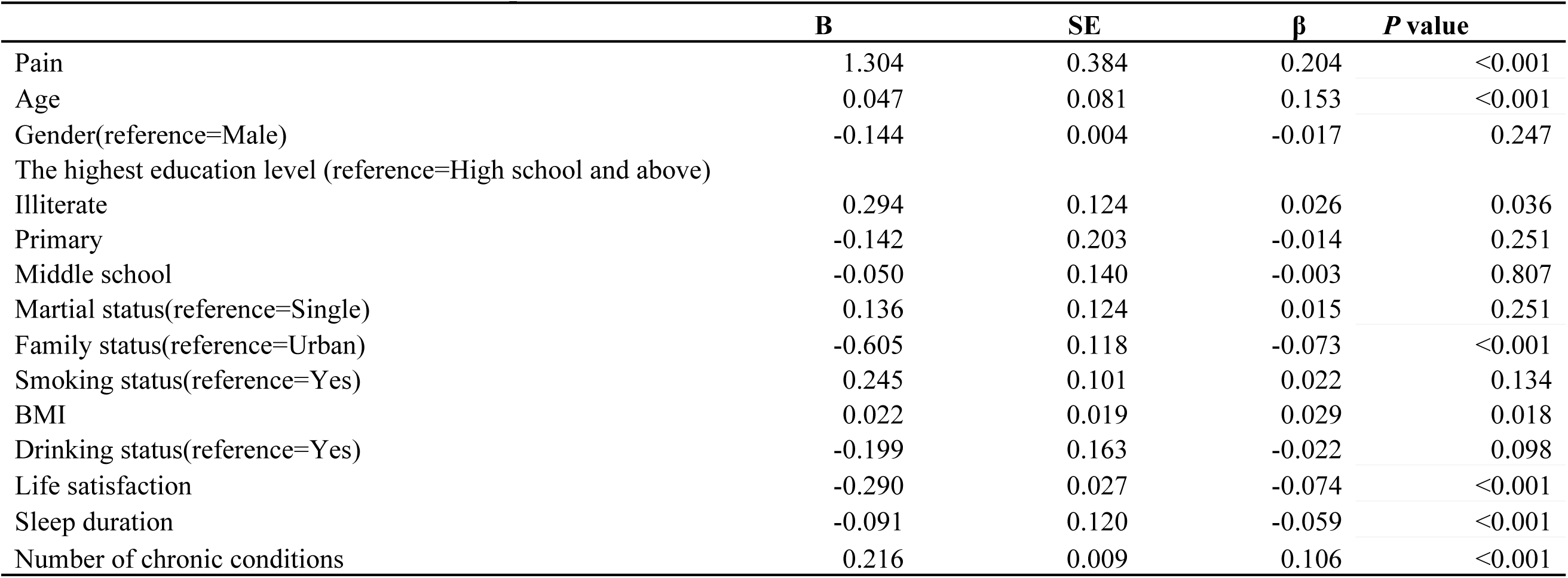
The association between pain and functional limitation.

**Table 4.**
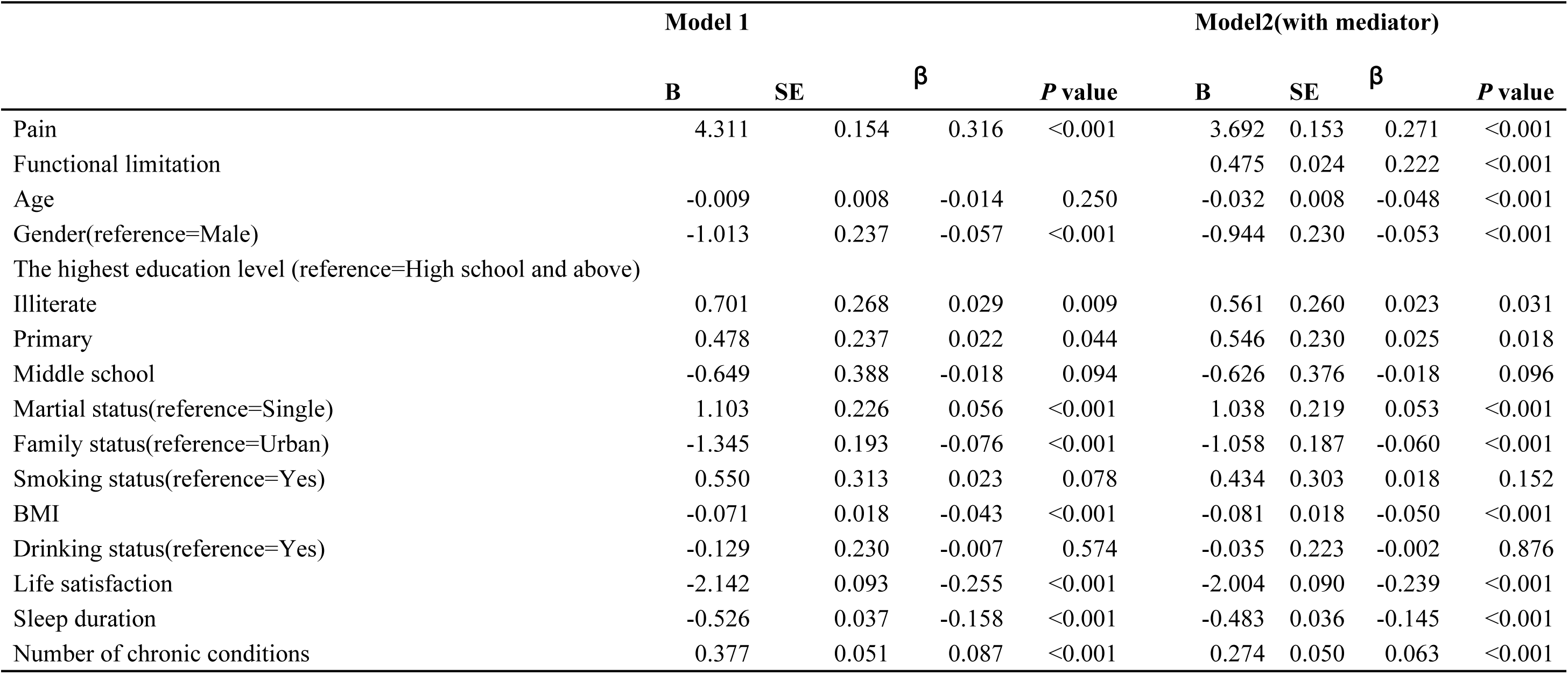
The association between pain and depression with functional limitation.

## 4. Discussion

The current study is the first to investigate the mediating role of functional limitation in the relationship between pain and depression among a population-based cohort of middle-aged and older adults in China. Our findings indicate that baseline pain is significantly associated with increased depressive symptoms. This relationship persists even after accounting for functional limitation, suggesting that functional limitation partially mediates the association between pain and depression. In line with our hypotheses, functional limitations played a partial mediating role in the relationship between pain and depression. Consistent with previous studies^28^, our study suggested that pain increased the risk of depression. Research has shown that the sensory pathways involved in body pain overlap with brain regions associated with mood regulation. These regions include the insular cortex, prefrontal cortex, anterior cingulate cortex, thalamus, hippocampus, and amygdala. This anatomical relationship provides a structural basis for the coexistence of pain and depression^29^. The inflammatory response has been shown to play a significant role in the development of both pain and depression. However, when pain is mediated by inflammatory mechanisms, it may exhibit a stronger association with depression^30^. This relationship is further supported by evidence from studies examining brain regions and neurological functional systems, which reveal a close correlation between pain and depression^31^. Specifically, chronic pain, driven by persistent inflammatory processes, has the potential to lead to the onset of depressive symptoms.

Our results are consistent with previous studies that pain is associated with increased odds of functional limitation^32^. Central sensitization amplifies pain signals within the nervous system, resulting in heightened sensitivity and avoidance of activities that may exacerbate pain, ultimately decreasing mobility and strength^33^. The fear-avoidance model also illustrated how fear of pain can lead to avoidance behaviors, creating a cycle of disability and functional limitation^34^. Similarly, we found a positive correlation between functional limitation and depression. Functional limitation may reduce the independence of middle-aged and older adults, making it difficult for them to complete daily activities such as self-care, household chores, and social activities. This reduced independence and mobility may lead to feelings of loneliness, helplessness, and decreased self-esteem, which are common manifestations of depressive symptoms. Previous studies have shown that high growth trajectories in ADL and IADL significantly increase the risk of depression in middle-aged and older populations^35^ which is consistent with our research.

Importantly, the results of this study also suggest that functional limitation mediates the association between pain and depression. Empirical evidence suggests that individuals experiencing chronic pain are more likely to report reduced physical functioning, which exacerbates emotional distress and increases the risk of depression^36^. This finding provides new evidence in the existing literature that pain may increase subsequent depression through functional limitation in middle-aged and older adults.

The results of this study have important theoretical implications for understanding the relationship between pain and depression. First, clinicians and mental health professionals should pay attention to the functional limitation of patients with pain as an important indicator of the risk of depression and intervention. Second, interventions that target functional limitation, such as rehabilitation, psychological support, and social engagement, may help alleviate depressive symptoms in patients with pain. Lastly, communities and medical institutions can carry out comprehensive intervention programs for patients with pain, aiming to improve their self-care and psychological resilience, thereby reducing the risk of depression.

There are some limitations to this study. Firstly, due to the cross-sectional nature of the data, it was not possible to establish a causal relationship between pain, functional limitation, and depression. Future studies should adopt a longitudinal design to better assess the dynamic relationship between these variables. Secondly, the measurement of functional limitation relies primarily on patient self-report and may be subject to recall bias or subjectivity. Future research could incorporate objective functional assessment tools to improve the accuracy of measurements. In addition, this study did not analyze the severity and duration of pain and functional limitation, which warrants further exploration in future studies. Finally, due to the limitations of sample size and follow-up time, the results of this study may not fully reflect the complex relationship between pain and depression. Therefore, caution should be exercised in interpreting the results of this study.

## 5. Conclusions

In conclusion, this study reveals the mediating role of functional limitation in the relationship between pain and depression, and emphasizes the importance of paying attention to functional limitation in patients with pain in clinical practice. Future studies should further validate this mediation mechanism and explore more potential interventions to improve the physical and mental health of patients with pain.

## Data availability statement

The datasets generated and/or analyzed for the current study are available in the CHARLS repository at http://charls.pku.edu.cn.

## Ethics statement

This study was based on publicly available datasets (CHARLS). Ethical review and approval were not required for the study on human participants in accordance with the local legislation and institutional requirements. Written informed consent from the participants or the participants’ legal guardian/next of kin was not required to participate in this study in accordance with the national legislation and the institutional requirements. CHARLS was approved by the Biomedical Ethics Review Committee of Peking University (IRB00001052-11015); the patients/participants provided their written informed consent to participate in this study.

## Author Contributions

FJ: Formal analysis, Writing – original draft. FNY: Funding acquisition, Supervision, Writing – original draft, Writing – review & editing.

## Generative AI statement

The authors declare that no Gen AI was used in the creation of this manuscript.

## Publisher’s note

All claims expressed in this article are solely those of the authors and do not necessarily represent those of their affiliated organizations, or those of the publisher, the editors and the reviewers. Any product that may be evaluated in this article, or claim that may be made by its manufacturer, is not guaranteed or endorsed by the publisher.

---

**S1 Fig.**
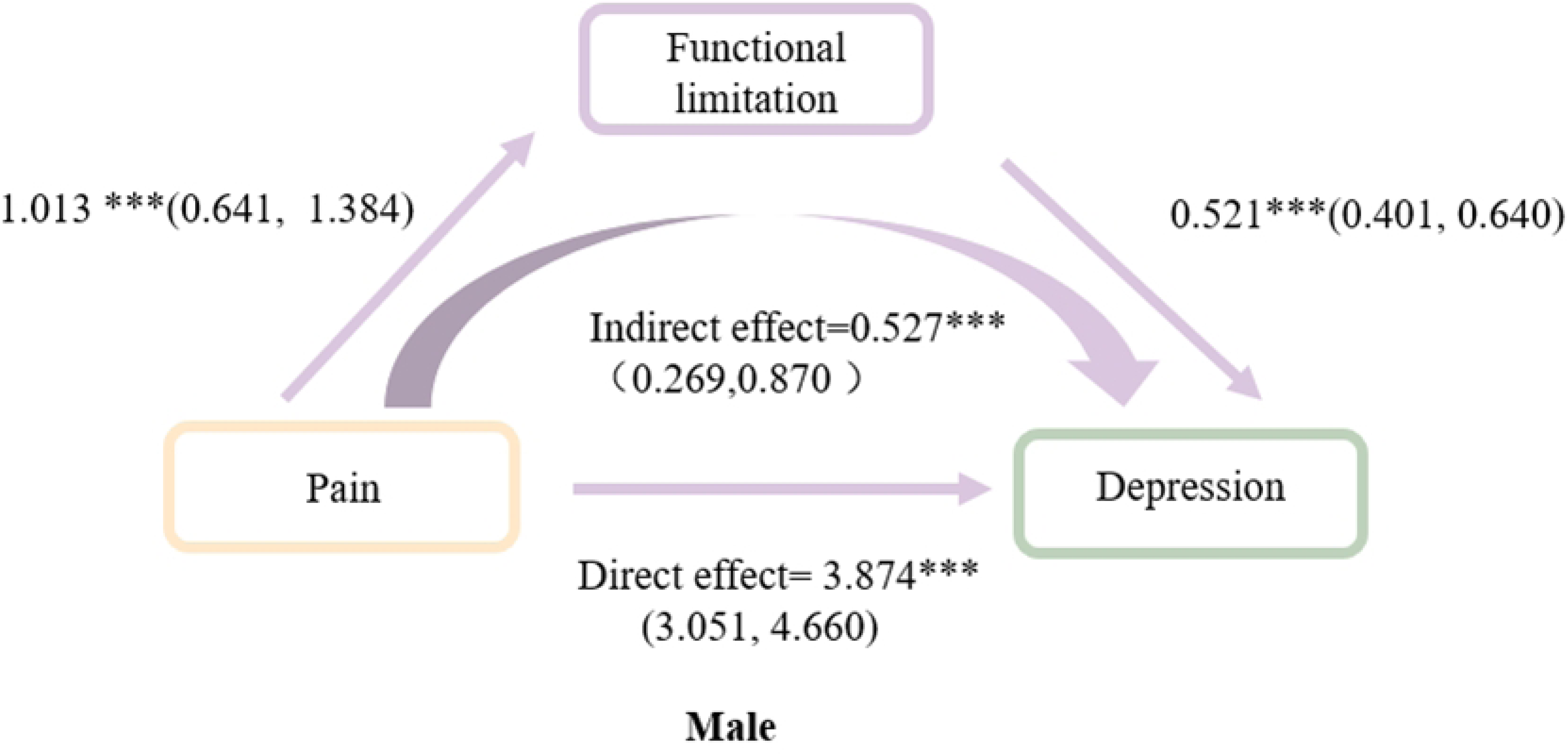
Mediation model illustrating the relationship between pain and depression mediated by functional limitation in male. Abbreviations: *, ***: significantly different from 0.05 to 0.001.

**S2 Fig.**
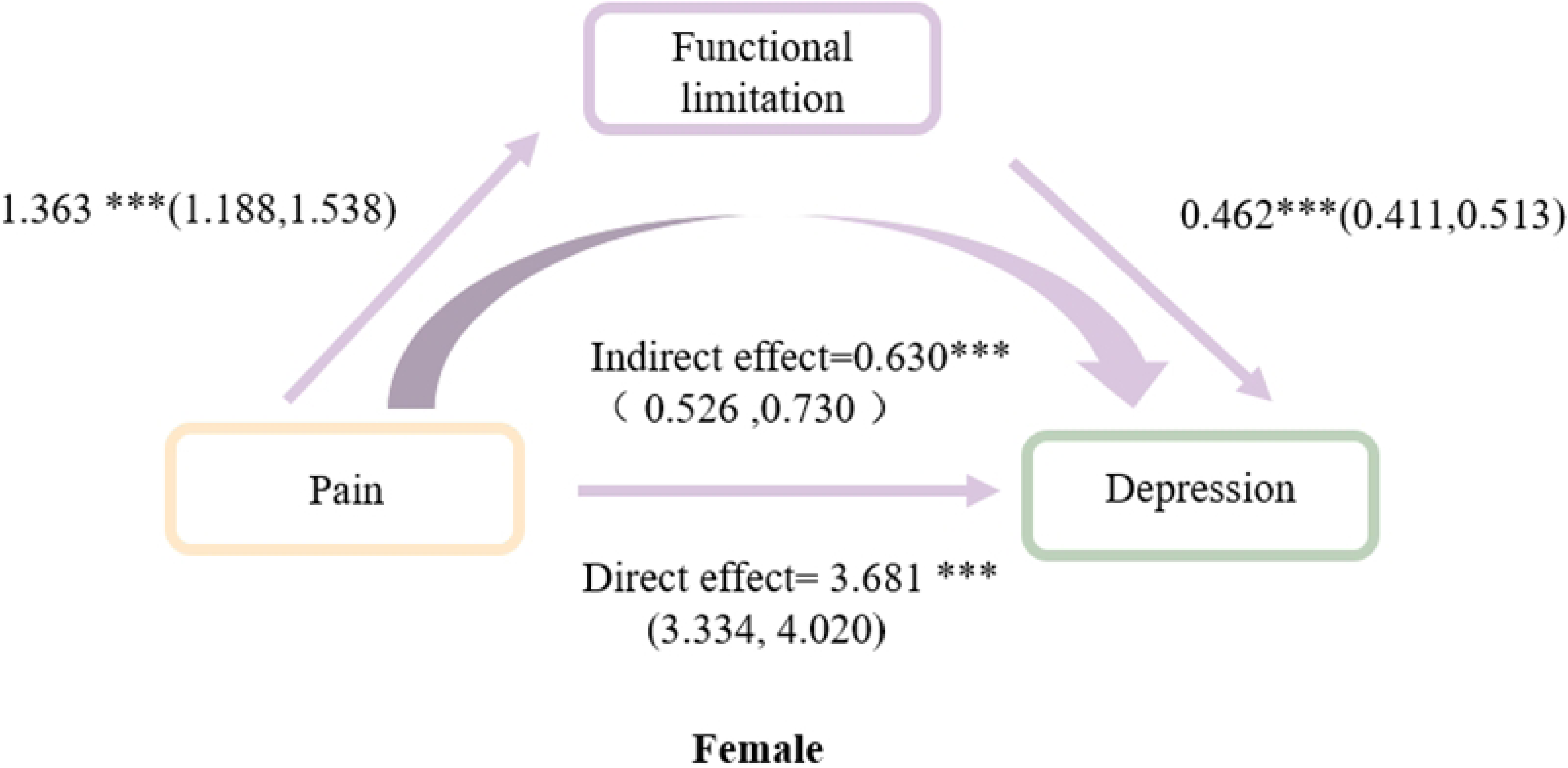
Mediation model illustrating the relationship between pain and depression mediated by functional limitation in Female. Abbreviations: *, ***: significantly different from 0.05 to 0.001.

**S3 Fig.**
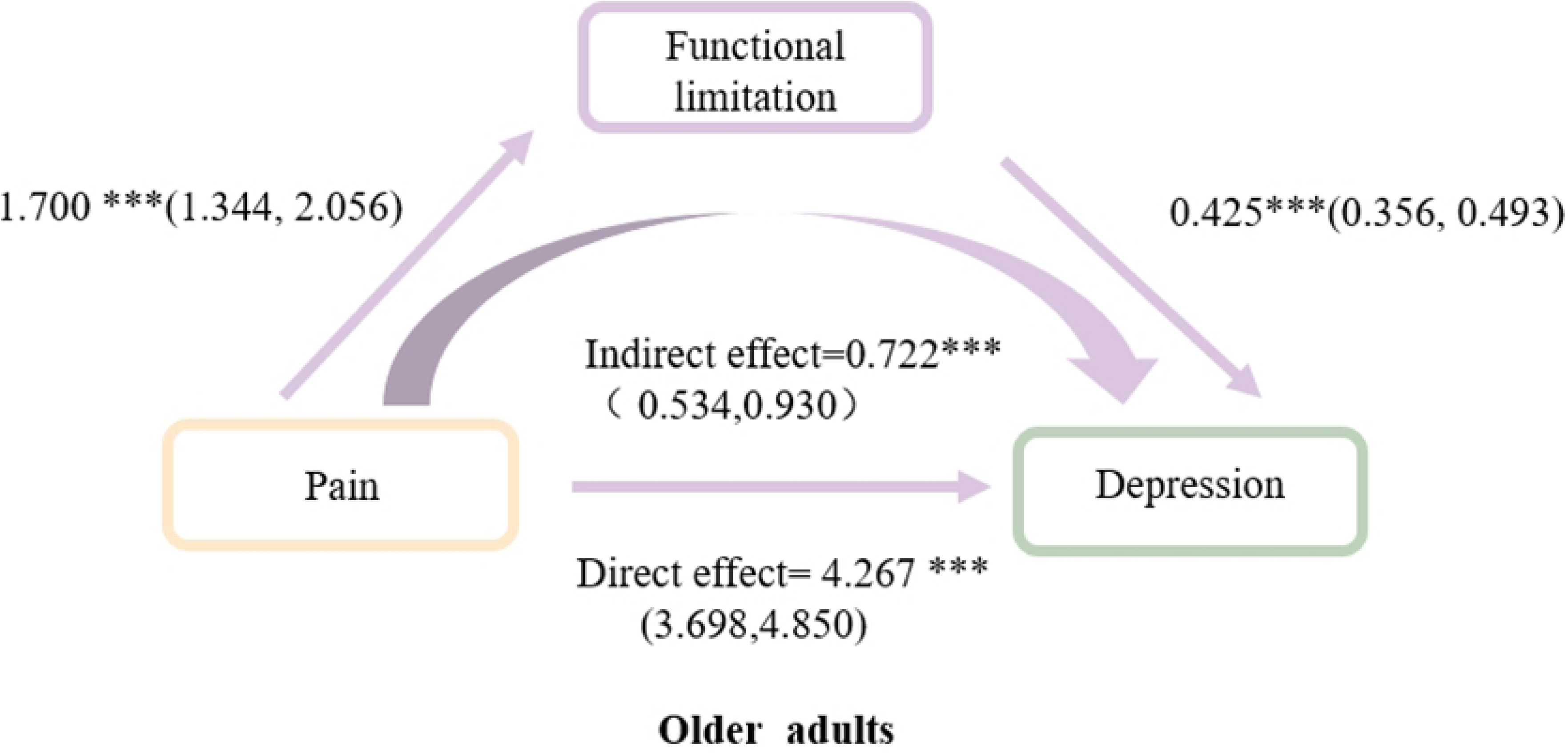
Mediation model illustrating the relationship between pain and depression mediated by functional limitation in older adults. Abbreviations: *, ***: significantly different from 0.05 to 0.001.

**S4 Fig.**
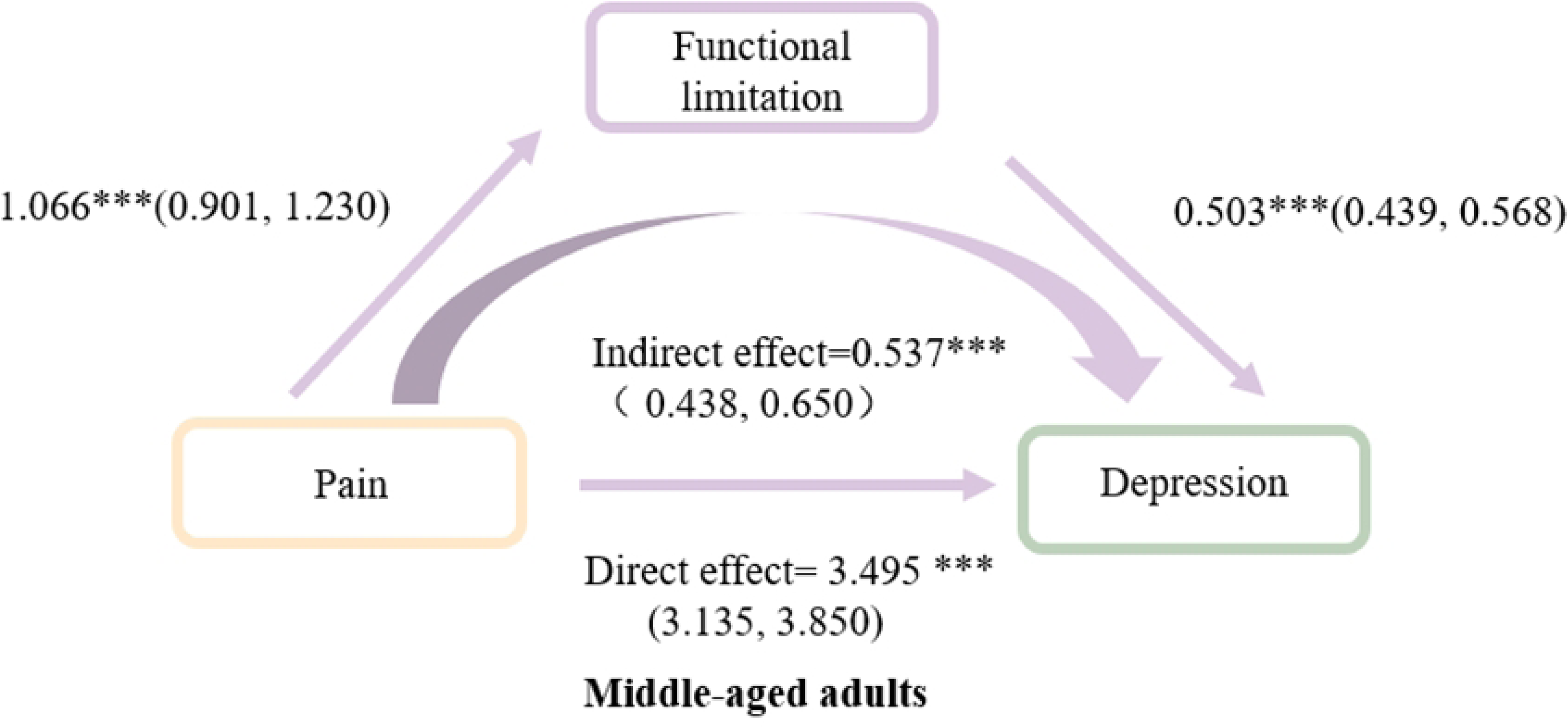
Mediation model illustrating the relationship between pain and depression mediated by functional limitation in middle-aged adults. **Abbreviations:** *, ***: significantly different from 0.05 to 0.001.

